# Lytic and temperate phage naturally coexist in a dynamic population model

**DOI:** 10.1101/2022.12.09.519798

**Authors:** Ofer Kimchi, Yigal Meir, Ned S. Wingreen

## Abstract

Obligate lytic and temperate phage preying on the same bacteria coexist despite the presumption that a single resource should only support a single competitor. We construct a mathematical model demonstrating that such coexistence is a natural outcome of chaotic dynamics arising from competition among multiple phage and their lysogens. While obligate lytic (virulent) phage populations typically dominate, surprisingly, they also more readily fluctuate to extinction within a local community.

---

Phage—viruses that infect bacteria—are subject to strong evolutionary pressures. One optimization axis is the lysis-lysogeny decision phage face when infecting a bacterium. Upon infection, phage can either lyse the bacterium, generating a burst of phage progeny, or lysogenize the bacterium, incorporating the phage genome into the bacterium. The resulting lysogen is generally immune to subsequent infection by the same class of phage [1, 2].

Some phage, such as Escherichia coli phage T4, never lysogenize their bacterial hosts, and are referred to as obligate lytic (or virulent) phage; others, such as the E. coli phages λ and P1 and the Vibrio cholerae phage CTX*ϕ*, can perform either lysis or lysogeny upon infection, and are referred to as temperate phage [3]. The optimal lysis/lysogeny tradeoff depends on environmental conditions [4, 5]: when susceptible bacteria are abundant, phage do better by lysing their hosts and releasing a burst of progeny; when these bacteria are scarce, phage are better off lysogenizing their hosts [6, 7]. However, lytic and temperate phage that prey on the same bacterial hosts are found to coexist with one another [8]. How can they coexist if one is more optimal than the other?

Explanations for the coexistence of competing species generally rely on the idea that different species have different ecological niches, e.g. in terms of resources or space [9, 10]. In contrast, here we show that naturally arising chaotic population dynamics are sufficient for the coexistence of obligate lytic and temperate phage.

We consider *N_c_* phage classes preying on a single bacterial species (Fig. 1). Bacteria grow at a rate *α*, and are limited by phage predation rather than resource limitation. Phage populations grow through lysis. Phage infect bacteria at a rate *k*, leading to either lysis or lysogeny. Phage strains within a class are distinguished by their (fixed) fraction of infections that lead to lysogeny, which we denote by *f*. If the phage performs lysis, it creates *b* new copies of the phage and kills the bacterium. Lysogens are immune to reinfection by a phage of the same class, and are spontaneously induced to undergo lysis at a rate *γ*. Phage die (or migrate away) at a rate *δ*, and phage that attempt to infect immune lysogens also die. Values for parameters are motivated by [11, 12] (see Supplementary Information Section S1). For simplicity, we assume attempts to form a double lysogen lead to death of the infecting phage. An extended model allowing for multiple lysogeny (Extended Data Fig. ED1) produces similar results; see Supplementary Information Section S2.

**Figure 1:**
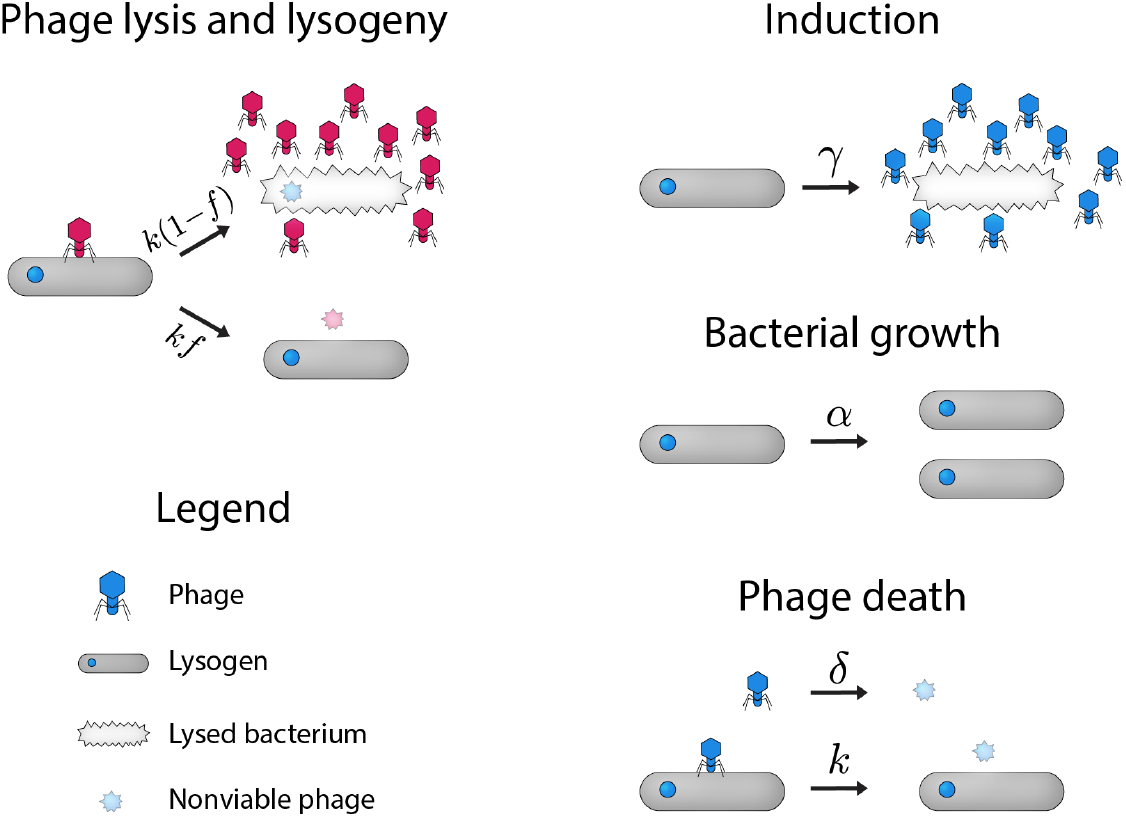
Model overview. A pictoral representation of the simplified model described by equations (1). Phage of different immunity classes are represented by different colors. Here, we disallow double lysogens and treat the population of sensitive (non-lysogenic) bacteria as negligible (see main text).

Each phage population *P*_*cf*_ is indexed by its immunity class *c* and strain *f*, i.e. the fraction of its infections leading to lysogeny. Strains with *f* = 0 are obligate lytic and have no associated lysogens. The populations of phage *P*_*cf*_ and their associated lysogens *L*_*cf*_ change according to:

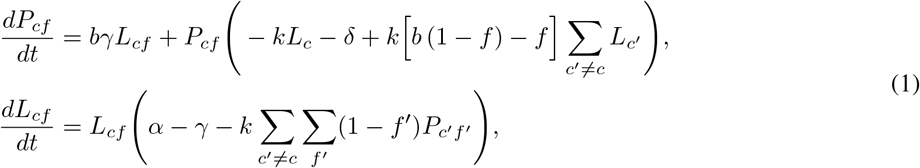

where *L*_*c*_ = ∑_*f*_ *L*_*cf*_. Equations (1) are a natural simplification of a more comprehensive model including both sensitive (i.e. non-lysogenic) bacteria as well as double lysogens and a finite lysis time; see Supplementary Information Section S2.

We find that when a single obligate lytic strain and a single temperate strain compete, one or the other goes extinct depending on whether they are of the same or different classes (Extended Data Fig. ED2). However, when multiple temperate strains compete, we observe sustained chaotic dynamics (Fig. 2a; Extended Data Fig. ED3a; Section S3 of Supplemental Information). We hypothesized that these frequent large variations in populations (or “boom and bust” cycles) could lead to an opportunity for obligate lytic phage.

**Figure 2:**
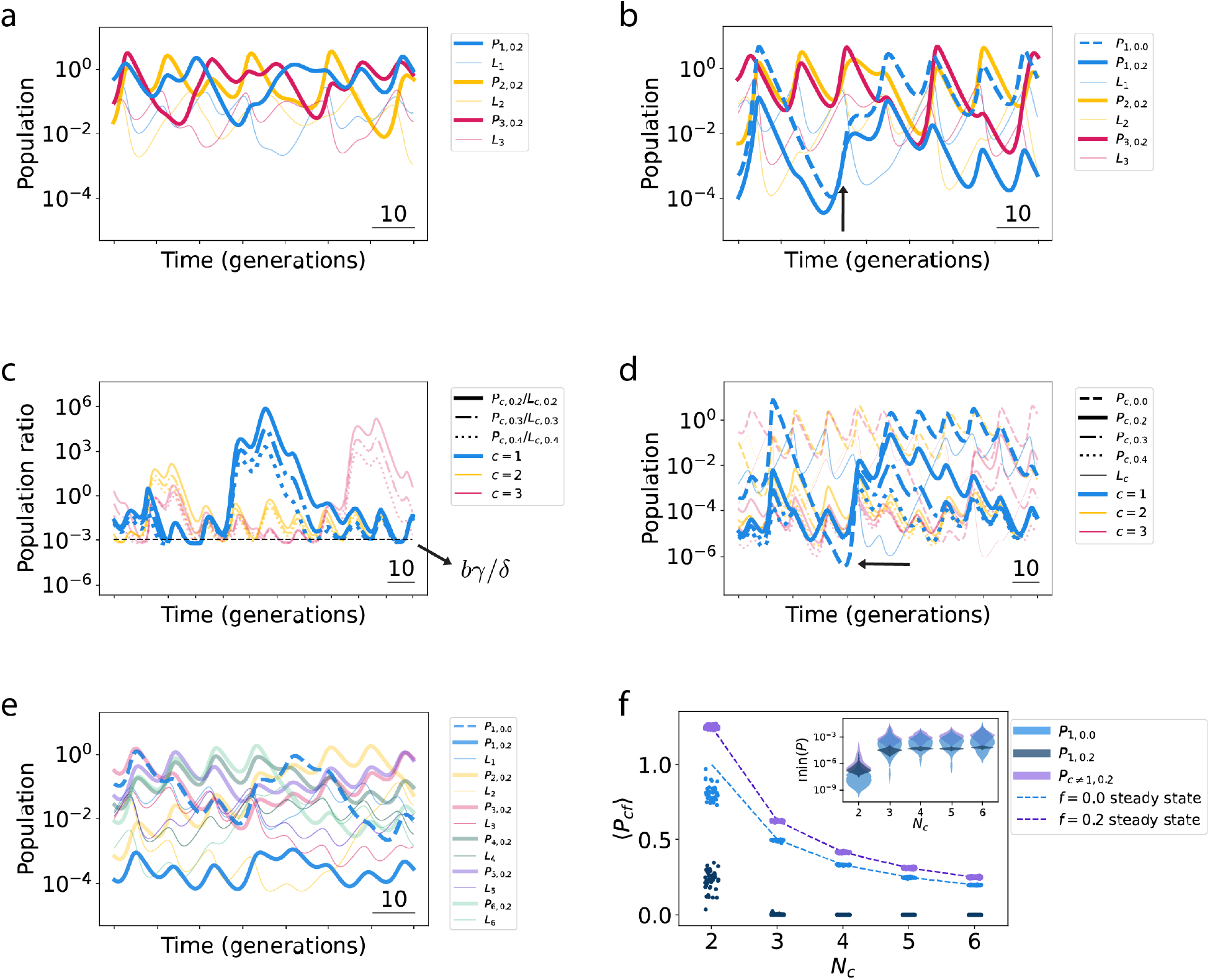
Population dynamics of coexisting obligate lytic and temperate phage. **a**, Competition among multiple temperate phage classes (*N*_*c*_ = 3) leads to chaotic dynamics. **b**, Chaos allows obligate lytic and temperate phage to coexist, with obligate lytic strains typically dominating over temperate strains of the same class. Arrow points to a growth period of phage of immunity class 1 (blue). **c**, Simulation of *N*_*c*_ = 3 phage immunity classes each with one obligate lytic strain and three temperate strains; all strains coexist together. Here, temperate phage populations are plotted normalized by their respective lysogen population. In periods of growth of a particular phage immunity class, the more lytic strains dominate, but phage-to-lysogen population ratios “bunch” together at troughs, with a population floor predicted by equation (2) (black dashed line). **d**, Plot of populations (rather than population ratios) of simulation shown in **(c)**. In order to compare the temperate phage population floor with a population dip of an obligate lytic phage which lacks such a floor (arrow), lysogens of different strains of the same class were initialized with equal populations (and remain equal indefinitely; see Extended Data Fig. ED4). **e**, Increasing the number of phage classes (here to *N*_*c*_ = 6) generally leads to smaller population variation. **f**, Summary statistics across 50 simulations, each for 2000 generations. Each simulation competes *N*_*c*_ − 1 immunity classes, each consisting of a single temperate strain (purple), along with a single class with two strains: one obligate lytic (light blue), one temperate (dark blue). The steady-state fixed point solutions for the phage populations (equation (3); dashed curves) agree well with the average of the chaotic trajectories. Average phage populations decrease with *N*_*c*_ (main figure, scatter plot) while minimum population sizes increase with *N*_*c*_ (inset, violin plot). Three simulations for *N*_*c*_ = 2 in which the obligate lytic phage fluctuated to extinction were excluded.

Obligate lytic strains are best able to capitalize on periods of phage population growth, since they turn all available susceptible bacterial hosts (i.e. lysogens from other immunity classes) into new phage. Thus, in conditions that allow for phage expansion, obligate lytic strains outcompete temperate strains of the same immunity class (Fig. 2b; arrow). Indeed, when an obligate lytic strain was introduced into the simulations of Fig. 2a, it typically dominated over the temperate strain of the same class (Fig. 2b). Surprisingly, both strains persisted at high population numbers. (The behavior of the phage in other immunity classes did not change qualitatively upon the introduction of the obligate lytic strain.)

How do temperate phage persist if obligate lytic phage outcompete them during periods of phage expansion? To probe this behavior further, we carried out simulations with three phage immunity classes, each with four strains (i.e. different values of *f*). We continued to see robust coexistence among all strains, and noticed a striking “bunching” effect when plotting the ratio of the temperate phage to their corresponding lysogen populations (Fig. 2c). During periods of growth of a particular phage class, strains with smaller lysogeny fractions *f* typically outcompete those with larger *f* ; however, *P/L* ratios were all nearly equal at the troughs (Fig. 2c). We traced this bunching effect to the induction of lysogens which buffers temperate phage populations against periods of decline. To test this explanation of the bunching phenomenon quantitatively, we consider the behavior of an obligate lysogenic (or “dormant”) phage with *f* = 1, whose dynamics are entirely determined by its respective lysogen. By setting *dP*_*c*1_*/dt* = 0, we find a homeostatic population for the phage, 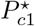, which is approximately proportional to the strain’s lysogen concentration:

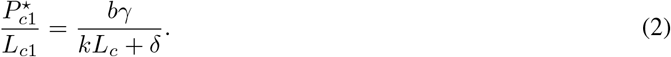

When the phage population dips below 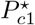, induction restores it back up, and when the phage population rises above 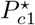, phage death pushes it down. As shown in Fig. 2c, equation (2) accurately predicts the population of temperate phage at the troughs. (We note that the simplified model of equations (1) retains a perfect memory of the initial ratios of lysogens of the same immunity class since *dL*_*cf*_ */dt* is independent of *f*. This feature is incidental to the observed bunching effect; see Extended Data Fig. ED4.) The bunching effect therefore implies a population “floor” for the temperate phage, protecting their populations during periods of decline.

In contrast, there is no population floor for the obligate lytic phage, as obligate lytic strains have no corresponding lysogens, and therefore no induction buffering their populations (Fig. 2d). Therefore, while obligate lytic strains typically outcompete temperate strains, lytic strain populations occasionally drop to very low levels. What then protects obligate lytic strains in nature from going extinct over long times?

Interestingly, in our model, the presence of more competitors leads to more stable behavior for all phage, raising the minima of obligate lytic strain populations. The reason for this stabilization is that the more phage classes present, the more distinct lysogenic species each one can prey on, and thus the smaller the fluctuations in the sum of the resources available to each phage (Fig. 2e-f). To explore this effect, we consider *N*_*c*_ − 1 classes each consisting of a single temperate strain, along with a single class with two strains (one obligate lytic, one temperate). Although the system dynamics are chaotic, the average population ⟨*P*_*cf*_⟩ of each phage with one strain in its immunity class agrees quantitatively with the steady-state fixed point value

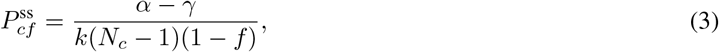

i.e. the non-zero solution to *dP/dt* = *dL/dt* = 0 (Fig. 2f). The average of the obligate lytic phage population is also reasonably well predicted by equation (3) with *f* = 0, despite the presence of a temperate strain of the same immunity class (Fig. 2f). The obligate lytic population is on average orders of magnitude larger than that of the temperate phage in its immunity class, but only slightly smaller than the populations of temperate phage of other immunity classes, as predicted by equation (3). Also as predicted by equation (3), as *N*_*c*_ is increased, the average population of each strain decreases; however, the minimum population of each strain *increases* and then plateaus with increasing *N*_*c*_ (Fig. 2f, inset). In fact, for *N*_*c*_ = 2, in 3 out of the 50 simulations, the obligate lytic strain fluctuated to extinction; no such extinction events occurred for *N*_*c*_ *>* 2. Thus, extinction becomes less likely in communities consisting of more phage immunity classes.

Our model makes three main simplifying approximations. First, while we considered the lysogeny fraction *f* for each strain to be a constant, as it is for, e.g., phage P1 [13], other phage such as *λ* vary *f* depending on the relative abundance of phage and bacteria [14, 15]. Second, we assume phage can only form single lysogens, an abridged version of more complex polylysogenic systems [2, 16]. Third, our model treats lysis times as negligible by considering small burst sizes, approximating the finite lysis times and larger burst sizes measured experimentally (see Section S1). While quantitatively significant, these approximations are not essential to our main results: an extension of our model to a more realistic system including sensitive bacteria and double lysogens yields the same qualitative behavior (Extended Data Fig. ED5), as do simulations including heterogeneous lysogeny probabilities (Extended Data Fig. ED3b-c).

Our model makes several experimentally-testable predictions. Following e.g. Ref. [2], a community of several phage immunity classes can be engineered such that the lysogens of each class can be infected by the other classes, but are immune to infection by strains within the same class. First, our model predicts that when a sufficient number of phage of different immunity classes (depending on the maximum number of cohabiting prophage within lysogens; see Extended Data Fig. ED5) are placed within a chemostat with a dilution rate slower than the death rate of lysogens, the populations within such a community will vary chaotically. Second, our results argue that introducing an obligate lytic phage in the same immunity class as one of the temperate phages will lead to a substantial (albeit finite) decrease of the latter’s population, with little effect on the behavior of the phage in other immunity classes. Third, our results predict that obligate lytic phage will have larger population fluctuations than temperate phage. Finally, we predict that while the average phage populations will decrease with increased number of immunity classes *N*_*c*_, the minimum phage populations taken over a long enough window will actually increase with *N*_*c*_.

Contrary to prior work finding optimal phage strategies survive while sub-optimal strategies go extinct [11, 17] our results suggest coexistence of different strategies may be commonplace. With a single obligate lytic phage strain and a single temperate phage strain of different immunity classes (i.e. a model akin to that considered in Extended Data Fig. ED2, bottom row), Stewart & Levin previously showed static steady-state coexistence is possible for certain parameter combinations, such as when the obligate lytic phage burst size and infection rate are smaller than those of the temperate phage [18]. Our work expands beyond these scenarios, finding for multiple classes a qualitatively different form of coexistence mediated by chaos, even when there is no difference between the parameters governing the behavior of the different phage strains. Species coexistence mediated by chaotic population dynamics has been modeled in other contexts [10, 19]; here, chaos arises naturally from the interactions of phage with their lysogens.

Our results suggest a natural bet-hedging mechanism for phage on the pan-genome level. When susceptible bacteria are plentiful, obligate lytic strains thrive, while their temperate cousins of the same immunity class persist at lower populations. When conditions worsen, however, temperate strains can outcompete obligate lytic strains. Thus, obligate lytic strains, typically dominant, would be first to go extinct. Here, the bet-hedging is not a product of the behavior of individual organisms, but rather a feature of competing strains within a larger, genetically related, population.

## Code Availability

All code used to generate the results and figures in this study can be found at https://github.com/ofer-kimchi/lytic-temperate-coexistence.

## Acknowledgements

We thank Jonathan Levine for useful discussions. This work was supported in part by the National Science Foundation through the Center for the Physics of Biological Function (PHY-1734030) and by the Peter B. Lewis ‘55 Lewis-Sigler Institute/Genomics Fund through the Lewis-Sigler Institute of Integrative Genomics at Princeton University (O.K.). This work was performed in part at Aspen Center for Physics, which is supported by National Science Foundation grant PHY-1607611.

## Author Contributions Statement

All the authors designed research, analyzed data, and wrote the article. O.K. wrote Python code.

