## Supplementary Information for "Lytic and temperate phage naturally coexist in a dynamic population model"

#### Extended data figures

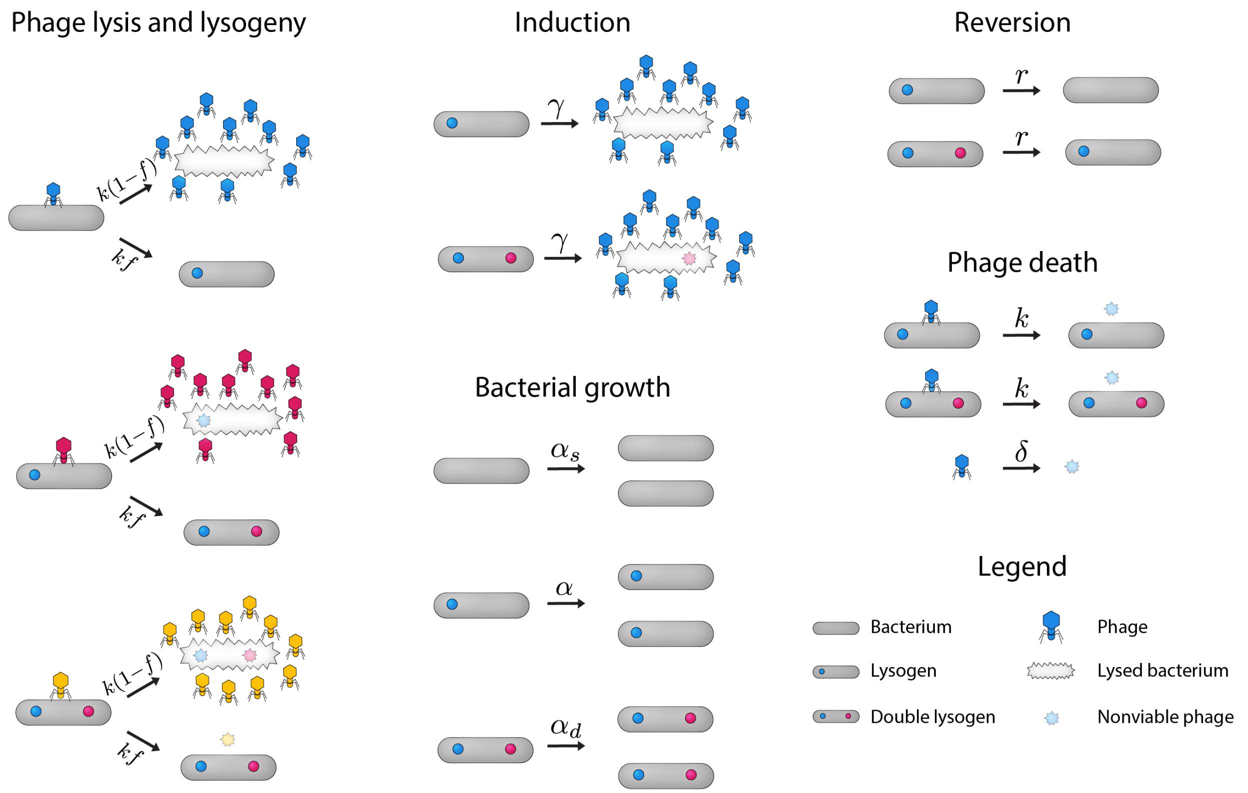


Figure ED1: **Comprehensive model overview.** A pictoral representation of the full model described by equations (S3). Phage of different immunity classes are represented by different colors.

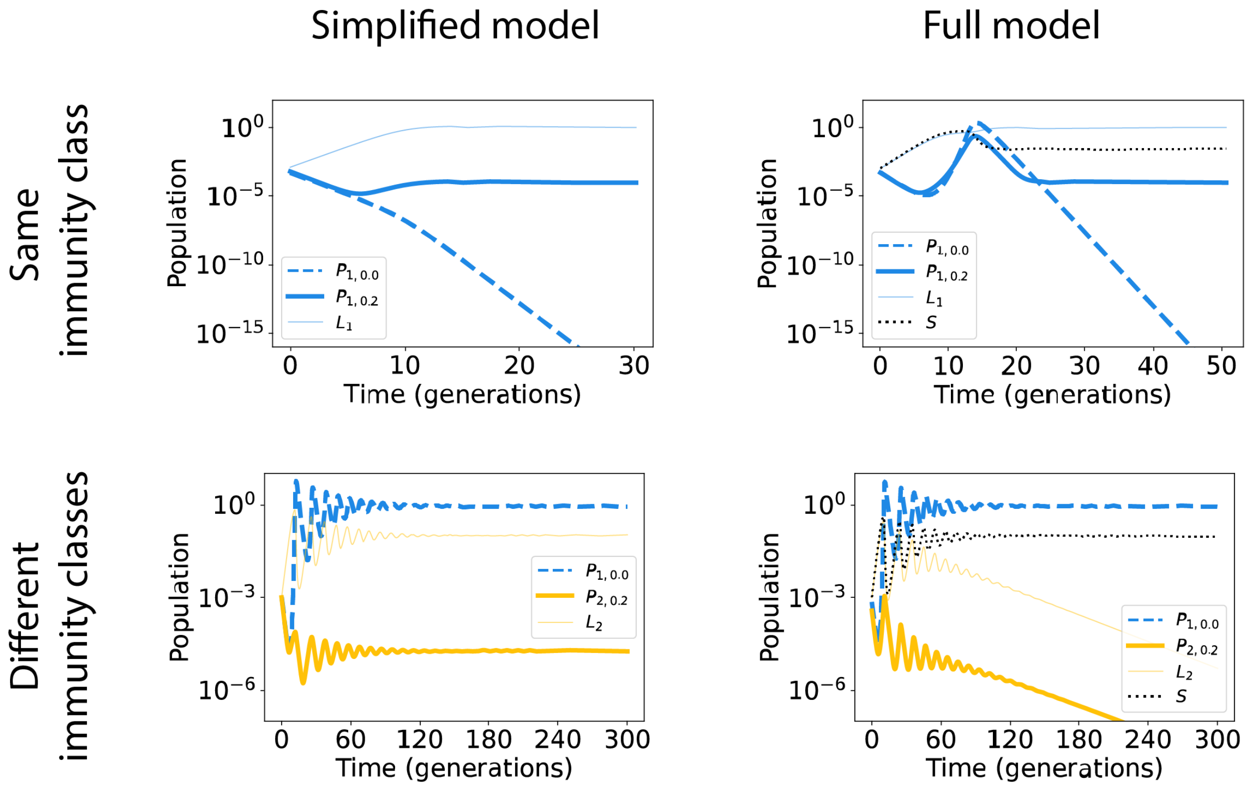


Figure ED2: **Competition between a single obligate lytic strain and a single temperate strain.** We show the results of competition between a single obligate lytic strain and a single temperate strain of either the same (top row) or different (bottom row) immunity classes. Both the simplified (left column) and more comprehensive (i.e. including sensitive bacteria, right column) models show that when the two strains are of the same immunity class, the obligate lytic strain goes extinct, while the temperate strain survives. When the two strains are of different immunity classes, the obligate lytic phage dominates over the temperate phage as the temperate-phage lysogens are not immune to the lytic phage. In the comprehensive model, the sensitive strain outcompetes the lysogens as the former has a slightly higher growth rate, leading the temperate strain in that model to go extinct; in the simplified model, lysogens survive and so the temperate strain persists as a result of induction. In equations (S3) and (S4) for the full model, the carrying capacity of bacteria was set to $K = 1$.

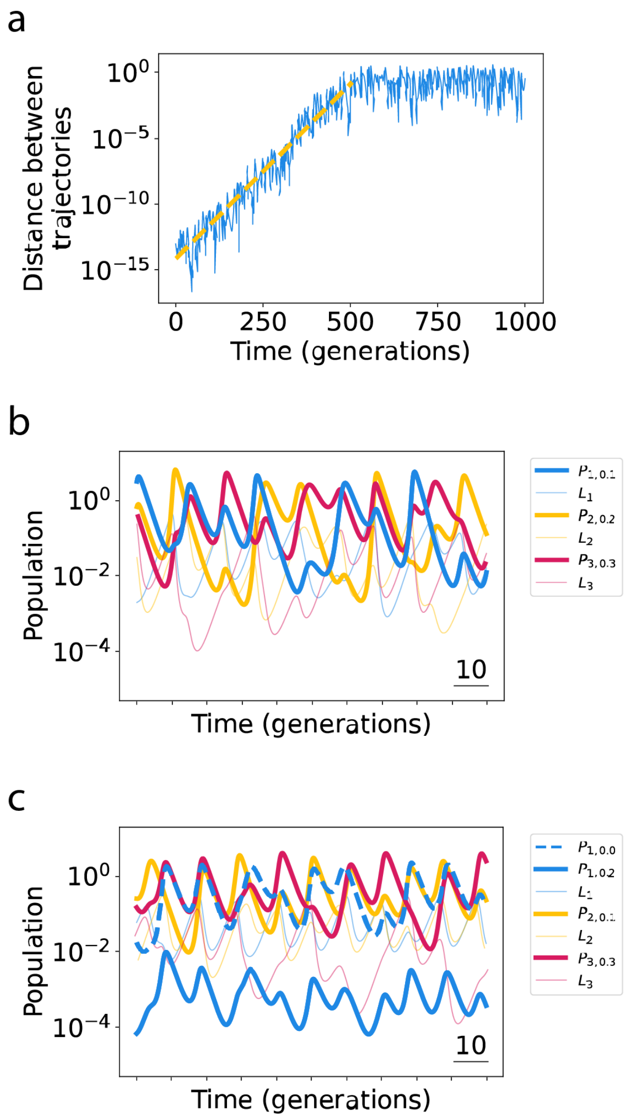


Figure ED3: **Trajectories display robust chaotic dynamics. a,** Three temperate phage of different immunity classes were simulated. Then, a second simulation with nearly identical initial conditions was run (initial phage populations were set to be $10^{-13}$ larger). We plot in blue the distance between the two trajectories of the first immunity class, $\sqrt{(P_{1,0.2} - P'_{1,0.2})^2}$. The initial portion of the plot is fit to an exponential, with Lyapunov exponent $9 \times 10^{-2}$ (yellow dashed line). That nearby trajectories diverge exponentially is a hallmark of chaotic dynamics. **b, c,** Chaotic dynamics persist with heterogeneous lysogeny fractions $f$, with little qualitative difference from the case of homogeneous $f$ (Fig. 2a-b).

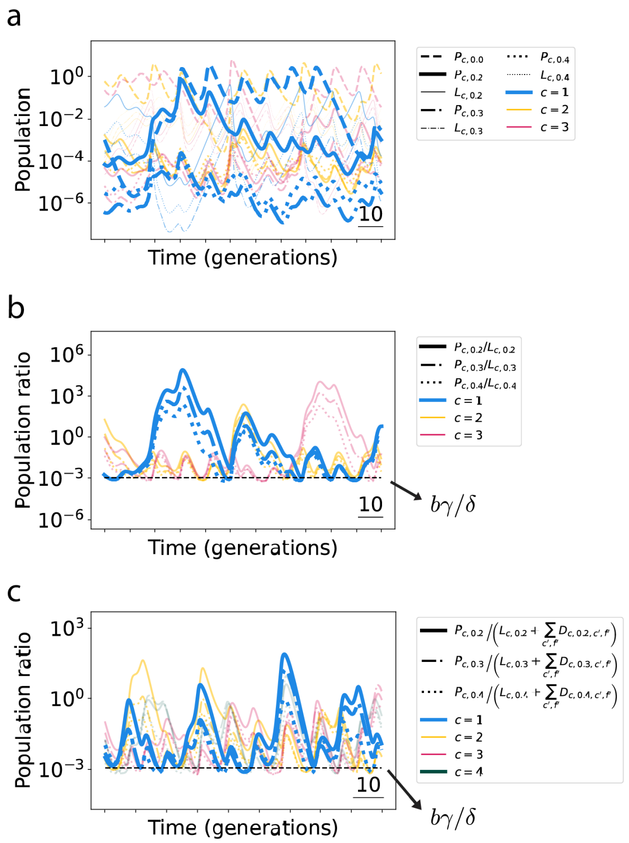


Figure ED4: **Bunching effect with heterogeneous initial lysogen populations.** **a,** Simulations performed as in Fig. 2c-d, i.e. with $N_c = 3$ phage immunity classes each with one obligate lytic strain and three temperate strains, but with one difference: here, initial lysogen populations were chosen randomly over a range of $\sim 2$ orders of magnitude. Since in the simplified model, equations (1), $dL_{c,f}/dt$ is independent of $f$, the lysogens of different strains of the same immunity class vary in lockstep together, and the system retains a perfect memory of their initial population ratios. **b,** The bunching effect at phage population troughs is independent of this memory, and the ratio of the population of each phage to the population of its respective lysogen is qualitatively unchanged compared to Fig. 2c. **c,** The full model (equations (S3)), which lacks this memory, displays the same bunching effect; furthermore, the population floor is quantitatively unchanged from the simplified model, and is determined by the total population of each phage strain's respective lysogens, including both single and double lysogens.

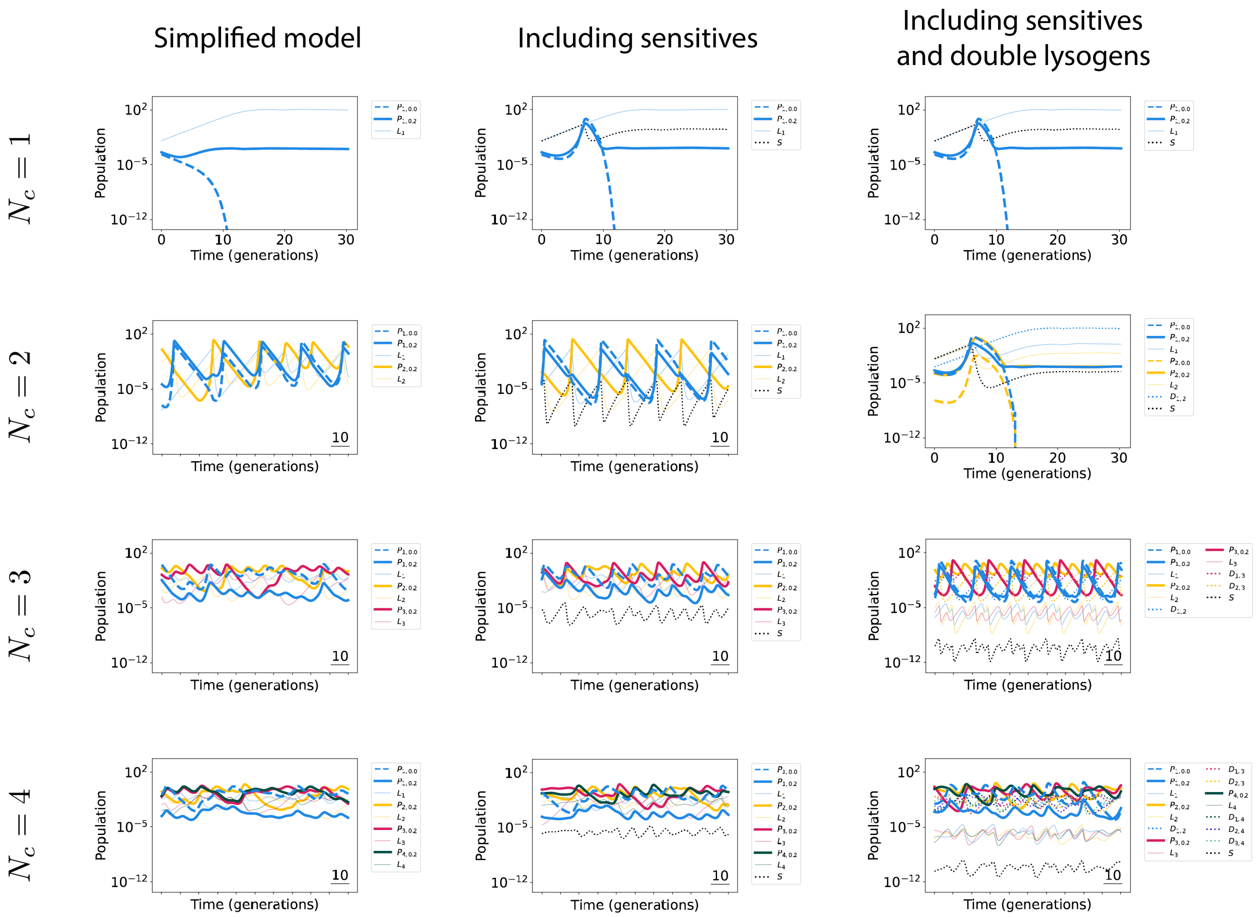


Figure ED5: **Comparison of models with different degrees of simplification.** Simulation results are shown for the simplified model (equations (1), left column), the model including sensitive (i.e. non-lysogenic) bacteria (equations (S4), middle column), and the full model including both sensitive bacteria and double lysogens (equations (S3), right column). Each simulation has $N_c$ competing phage immunity classes. One immunity class has both an obligate lytic strain and a temperate strain, while the other $N_c - 1$ immunity classes have a single temperate strain. Carrying capacity of bacteria, $K$, was set to 100 in the middle and right columns.

### S1 Parameters

The values for the parameters used in the model are motivated by Cortes *et al.* (2019) [11]. Parameters setting the units for time and population are the lysogen growth rate $\alpha$ and the infection rate constant $k$, which are both set to unity. The induction rate is small compared to the lysogen growth rate: $\gamma = 10^{-4}\alpha$. With this value of $\gamma$, induction of one lysogenic offspring will occur roughly 12 generations after lysogeny.

Our parameter choices differ from those of Cortes *et al.* in three main ways. First, while we consider finite nutrient conditions (and thus finite bacterial carrying capacities) in Section S2, we simplify the model in the main text to assume bacterial populations are exclusively limited by phage predation. Second, we assume a higher phage degradation rate than Cortes *et al.* by setting $\delta = \alpha$, accounting for both phage degradation and migration out of the local environment.

The third way our parameter choices differ from those of Cortes *et al.* regards the lysis time. Once a phage decides to undergo lysis, it creates many new phage virions. This process takes a significant amount of time, of the same order as a typical bacterial generation time. For example, phage $\lambda$ has a period of $\sim 50$ minutes between infection and lysis of its *E. coli* host, roughly 2.5 times as long as the doubling time of *E. coli* in the same experiment (20 minutes) [12]. Below, we describe how a model with zero delay and a small burst size leads to the same overall phage growth rate as a model with a finite delay and larger burst size.

To model a finite lysis time explicitly, we could in principle modify equation (1) into a delay differential equation with a delay time $\tau$:

$$\frac{dP_{c0}(t)}{dt} = kb_\tau P_{c0}(t-\tau)\sum_{c'\neq c} L_{c'}(t-\tau) - \delta P_{c0}(t), \tag{S1}$$

where for simplicity, we have considered the obligate lytic phage ($f = 0$), and assumed it is the only strain in immunity class $c$. We have denoted the burst size in this model by $b_\tau$ to distinguish it from the burst size in the model with $\tau = 0$. Also for simplicity, we approximate the total lysogen population susceptible to our strain of interest by a constant, $L$. We then have:

$$\frac{dP_{c0}(t)}{dt} = kb_\tau L P_{c0}(t-\tau) - \delta P_{c0}(t). \tag{S2}$$

The solution to this equation approaches an exponential for large $t$. Keeping all other parameters equal (here we use $L = 0.2$), we find that the solution for finite $\tau$ is well-approximated by a model with $\tau = 0$ and a smaller burst size $b \equiv b_0$. The dependence of $b$ on the time-delayed burst size $b_\tau$ and the length of the delay $\tau$ is shown in Fig. S1a. For realistic parameters, $\tau = 51$ min and $b_\tau = 170$ [12], we obtain an equivalent model with $\tau = 0$ and burst size $b = 12$ (rounded down from a best-fit value of 12.5). This simplified model with $\tau = 0$ gives a very good approximation to the results with a finite value of $\tau$ and a larger burst size, as shown in Fig. S1b.

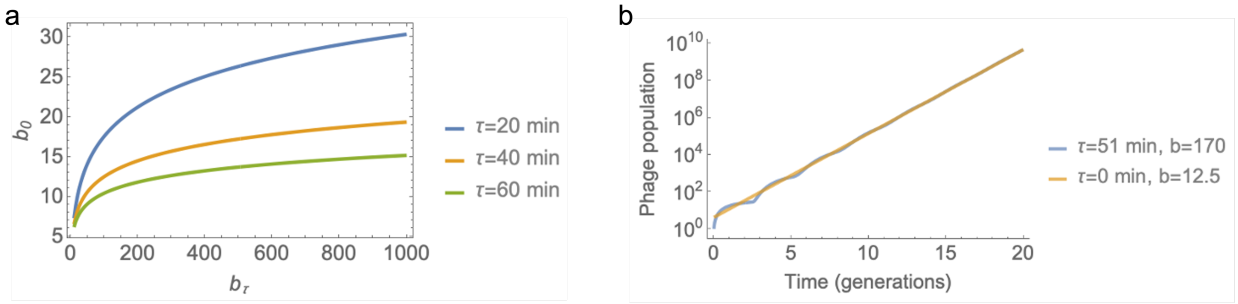


Figure S1: **A model with zero lysis time and a small burst size quantitatively approximates one with a finite lysis time and a larger burst size. a,** As a function of the true burst size $b_\tau$ for a given lysis time $\tau$, the best-fit burst size for a model with zero lysis time, $b_0$, is shown. **b,** The solution to the delay-differential equation with $\tau = 51$ min and $b_\tau = 170$ is plotted alongsize the solution with $\tau = 0$ and $b_0 = 12.5$.

### S2 More comprehensive model

The model considered in the main text is a natural simplification of a more comprehensive model including both sensitive (i.e. non-lysogenic) bacteria and double lysogens. In this section, we discuss this more comprehensive approach. Main qualitative results are unchanged between the models (Fig. ED5).

This model is summarized in Fig. ED1, which pictorally depicts the following set of equations:

$$
\begin{aligned}
\kappa =& 1 - \Big(S + \sum_c L_c + \frac{1}{2}\sum_{c,c'} D_{c,c'}\Big)/K, \\
\frac{dS}{dt} =& \alpha_S \kappa S - kS\sum_c P_c + r\sum_c L_c, \\
\frac{dL_{cf}}{dt} =& \alpha\kappa L_{cf} + kfSP_{cf} - (\gamma + r)L_{cf} - kL_{cf}\sum_{c'\neq c} P_{c'} + r\sum_{c'} D_{cf,c'}, \\
\frac{dD_{cf,c'f'}}{dt} =& \alpha_D \kappa D_{cf,c'f'} + kfL_{c'f'}P_{cf} + kf'L_{cf}P_{c'f'} - D_{cf,c'f'}\Big[2(\gamma + r) + k\sum_{c''\neq c,c'}\sum_{f''}(1-f'')P_{c''f''}\Big], \\
\frac{dP_{cf}}{dt} =& b\gamma\Big(L_{cf} + \sum_{c'} D_{cf,c'}\Big) + \\
& P_{cf}\left(-k(L_c + \sum_{c'} D_{c,c'}) - \delta + k\Big[b\,(1-f) - f\Big]\Big[S + \sum_{c'\neq c}(L_{c'} + \frac{1}{2}\sum_{c''\neq c} D_{c',c''})\Big]\right),
\end{aligned} \tag{S3}
$$

where we have defined a reversion rate $r$ for a lysogen to lose a prophage (e.g. via mutation), a carrying capacity $K$ for bacteria (which implies an associated population-dependent growth rate modifier $\kappa$), and different growth rates $\alpha_S$, $\alpha$, and $\alpha_D$ for the sensitive bacteria $S$, single lysogens $L$, and double lysogens $D$, respectively. The double lysogens are defined such that $D_{cf,c'f'}$ is the population of double lysogens created by the infection of a sensitive bacterium by both $P_{cf}$ and $P_{c'f'}$. Infection can occur in any order, such that $D_{c,c'} = D_{c',c}$ (and $D_{c,c} = 0$). As previously, we have omitted the subscript $f$ to denote summing over the different strains of a given immunity class.

We assume that lysogens incur a small growth rate penalty, setting $\alpha_S = 1.05$ and $\alpha_D = 0.95$ (with $\alpha = 1$) [11]. We set the reversion rate $r$ equal to the induction rate $\gamma$. The results of simulations of these full equations for different values of $N_c$, with $K = 100$ (simulating non-limiting nutrient conditions), are shown in Fig. ED5 (right panel). In these simulations, we set one immunity class to have both obligate lytic and temperate strains, and the other $N_c - 1$ immunity classes to only have a single temperate strain.

We note that this comprehensive model lacks the perfect memory of initial lysogen populations implicit in the simplified model (equations (1)).

To recover the main-text equations from equations (S3), we make three approximations. The first of these is assuming that $D = 0$ for all strains, leading to the following set of equations (depicted in Fig. S2):

$$
\begin{aligned}
\kappa =& 1 - \Big(S + \sum_c L_c\Big)/K, \\
\frac{dS}{dt} =& \alpha_S \kappa S - kS\sum_c P_c + r\sum_c L_c, \\
\frac{dL_{cf}}{dt} =& \alpha\kappa L_{cf} + kfSP_{cf} - (\gamma + r)L_{cf} - kL_{cf}\sum_{c'\neq c} P_{c'}, \\
\frac{dP_{cf}}{dt} =& b\gamma L_{cf} + P_{cf}\left(-kL_c - \delta + k\Big[b\,(1-f) - f\Big]\Big[S + \sum_{c'\neq c} L_{c'}\Big]\right).
\end{aligned} \tag{S4}
$$

Results of simulations of these equations can be seen in Fig. ED5 (middle panel). This simplification yields the same qualitative picture as the full model, with one major exception: in the full model (including double lysogens), $N_c = 2$ behaves like $N_c = 1$ in the single-lysogen model. In both cases, a single bacterial strain immune to all phage takes

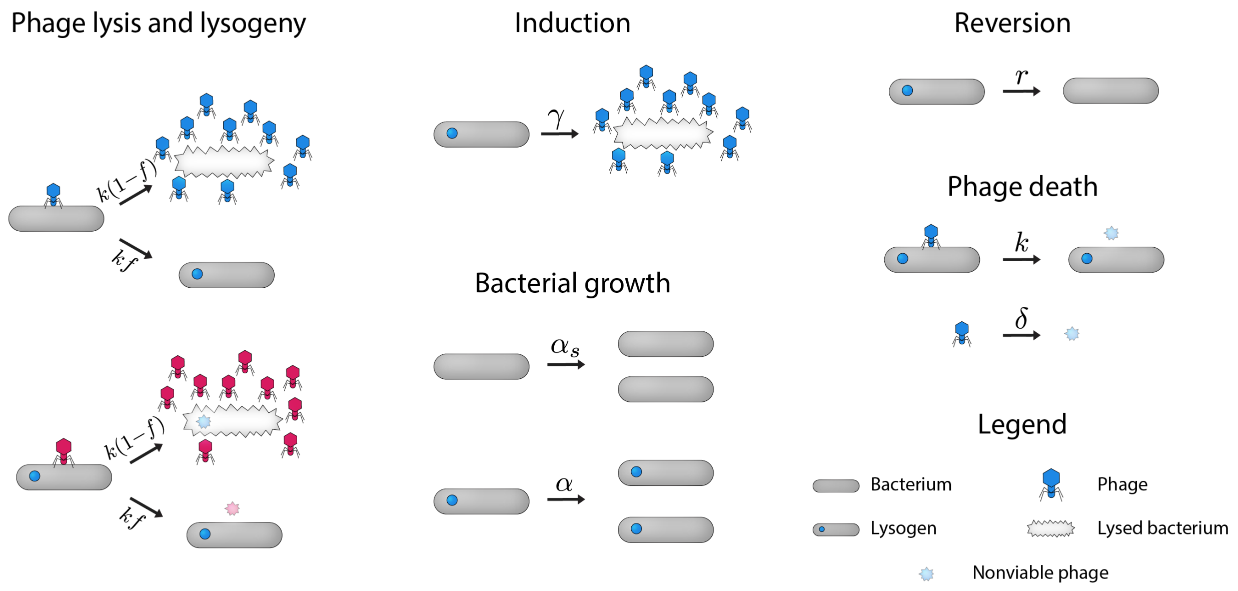


Figure S2: **Overview of model including sensitive bacteria.** A pictoral representation of the model described by equations (S4). Phage of different immunity classes are represented by different colors.

over the bacterial population, and leads to the extinction of obligate lytic phage strains, while temperate strains survive as a result of induction. Furthermore, $N_c = 3$ in the double-lysogen model behaves qualitatively like $N_c = 2$ in the single-lysogen model: both sometimes show oscillatory or quasi-oscillatory behavior. Finally, $N_c = 4$ in the double-lysogen model is qualitatively akin to $N_c = 3$ in the single-lysogen model: both display chaotic behavior. Thus, the model system displays chaotic behavior when there are at least two more immunity classes than the maximum allowed number of cohabiting prophage within lysogens. More generally, we find that the full model with $N_c$ immunity classes behaves similarly to the model disallowing double lysogens with $N_c - 1$ classes.

The other two approximations made to the full model to recover the main-text equations are to: 1) assume infinite $K$ (such that $\kappa = 1$), and 2) to set $S = 0$ (assuming zero reversion). Both are motivated by the results shown in Fig. ED5, which demonstrate that: 1) predation by phage limits bacterial populations, and 2) the population of sensitive bacteria is negligible, especially for $N_c > 2$. As shown in Fig. ED5 (left panel), these simplifications do not affect the qualitative behavior of the system.

#### S3 Chaotic behavior

Chaotic population trajectories are the predominant behavior found in the simplified model for $N_c \geq 2$, though not the only possibility: both a static fixed point and periodic oscillations sometimes occur. However, the latter have very small basins of attraction. We did not observe convergence to a fixed point in any of our simulations, but the fixed point can be solved for analytically. While simulations starting at the fixed point remain there, those that start even 1-5% away from it display chaotic behavior.

Some trajectories with $N_c = 2$ enter an oscillatory regime (as discussed above). We have also found some trajectories with $N_c = 3$, with a single temperate strain of $f = 0.2$ in each immunity class, that converge to an oscillatory regime after $10^4 - 10^5$ generations. In this case, a sudden variation of 50% in the populations returns the system to the chaotic regime. We have never observed periodic oscillations upon introducing an obligate lytic strain to these $N_c = 3$ simulations (Fig. S3a) nor do any of the trajectories in Fig. 2f show evidence of periodic oscillations. Similarly, we never observed periodic oscillations upon introducing a fourth identical phage immunity class (Fig. S3b), nor with $N_c = 3$ immunity classes where each strain has a different lysogeny fraction (Fig. S3c).

Therefore, given that natural variations in environmental conditions would frequently induce variability into the system, and given the natural heterogeneity among different phage populations, we do not expect periodic oscillations to be biologically relevant for $N_c > 2$.

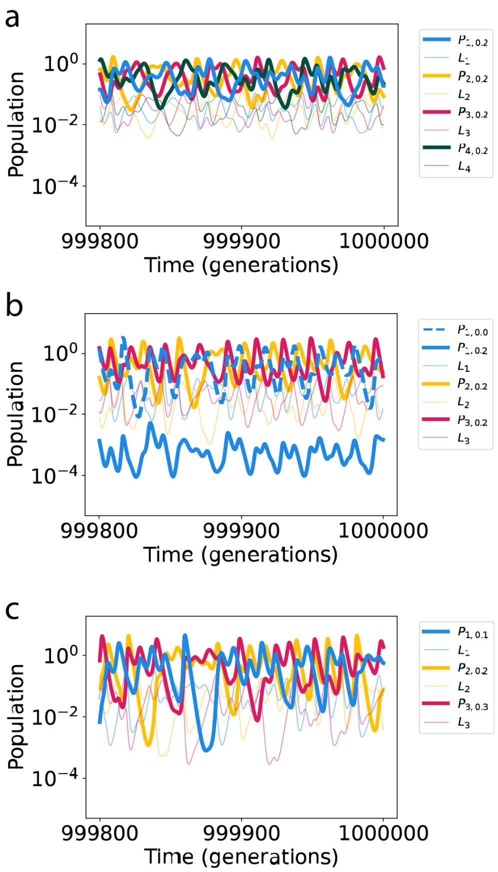


Figure S3: **Chaotic trajectories are typical. a,** Simulation with $N_c = 4$ phage immunity classes, each with a single temperate strain with lysogeny fraction $f = 0.2$. **b,** Simulation with $N_c = 3$ phage immunity classes, one of which has two strains—one obligate lytic and one temperate ($f = 0.2$)— and two of which have a single temperate strain ($f = 0.2$). **c,** Simulation with $N_c = 3$ phage immunity classes, each with a single temperate strain of different lysogeny fractions.
